# Haemocytes are critical for *Drosophila melanogaster* post-embryonic development, independent of control of the microbiota

**DOI:** 10.1101/2021.10.21.465347

**Authors:** HN Stephenson, R Streeck, A Herzig

**Affiliations:** Department of Cellular Microbiology, Max Planck Institute for Infection Biology, Charitéplatz 1, Berlin 10117

## Abstract

Proven roles for haemocytes (blood cells) have expanded beyond the control of infections in *Drosophila*. Despite this, the critical role of haemocytes in post-embryonic development has long been thought to be limited to control of microorganisms during metamorphosis. This has previously been shown by rescue of adult development in haemocyte-ablation models under germ-free conditions. Here we show that haemocytes have a critical role in post-embryonic development beyond their ability to control the microbiota. Using a newly generated, strong haemocyte-specific driver line for the GAL4/UAS system, we show that specific ablation of haemocytes is pupal lethal, even under axenic conditions. Genetic rescue experiments prove that this is a haemocyte-specific phenomena. RNA-seq data suggests that dysregulation of the midgut is a critical consequence of haemocyte ablation. We believe this novel role of haemocytes during metamorphosis is a major finding for the field. This is an exciting new *Drosophila* model to study the precise mechanisms in which haemocytes regulate tissue development, findings from which could have far reaching implications beyond invertebrate biology.

**Summary Statement:** Haemocyte-ablation in *Drosophila melanogaster* with a strong haemocyte-specific driver causes pupal lethality

## Introduction

*Drosophila melanogaster* is an important model to study both the immune and non-immune related functions of blood cells. There are 3 main blood cell types (haemocyte) in the fly (Mase, Augsburger et al. 2021). Plasmatocytes are macrophage-like cells, making up ∼95% of larval blood cell counts. In addition to apoptotic cell and microorganism phagocytosis, they secrete signalling peptides, anti-microbial peptides, and extra-cellular matrix (ECM) proteins (Braun, Hoffmann et al. 1998, Olofsson and Page 2005). Crystal cells account for ∼5% of larval blood cells. They express high levels of prophenoloxidases, which catalyse the extracellular production of melanin and toxic by-products upon cell lysis; critical for wound closure and immunity to a variety of pathogens (Binggeli, Neyen et al. 2014). Lamellocytes, rarely found in healthy larvae, transdifferentiate in large numbers from plasmatocytes to encapsulate large pathogens, such as wasp eggs (Sinenko, Shim et al. 2011). Recent single-cell RNA sequencing studies have shown greater heterogeneity in these cell types (Cattenoz, Sakr et al. 2020, Tattikota, Cho et al. 2020). There are ∼10 different sub-types of plasmatocytes that are differentiated by distinct processes, for example phagocytosis versus AMP production.

Two waves of haematopoesis occur in *Drosophila* development. Embryonic haemocytes originate from the head mesoderm; they are long-lived, many surviving until the adult stage (Tepass, Fessler et al. 1994). Larval haematopoesis occurs in the lymph gland and in haematopoeitic pockets, sessile patches of haemocytes associated with the larval cuticle. Haematopoetic pockets are the main source of increasing numbers of circulating haemocytes during larval development (Leitao and Sucena 2015); whereas haemocytes from the lymph gland are released into circulation at early metamorphosis (Jung, Evans et al. 2005)

Genetic ablation studies that aimed to identify the importance of blood cells for immune and non-immune related functions in *Drosophila* were first performed over a decade ago (Charroux and Royet 2009, Defaye, Evans et al. 2009, Shia, Glittenberg et al. 2009, Nehme, Quintin et al. 2011, Arefin, Kucerova et al. 2015). Haemocyte-specific expression of pro-apoptotic transgenes ablated cells through programmed cell death, achieving 60-75% reduction in larval haemocyte numbers. These studies primarily utilised promoters of the *Hemolectin* (*Hml*) gene, which shows haemocyte-specific expression in both larvae and adults (Sinenko and Mathey-Prevot 2004). Multiple studies showed a reduction in eclosion of adult flies of up to 60%; interestingly however, eclosion rates were rescued when larvae were reared with antibiotics or under germ-free conditions (Charroux and Royet 2009, Defaye, Evans et al. 2009, Shia, Glittenberg et al. 2009, Arefin, Kucerova et al. 2015). This suggested control of microorganisms by haemocytes is critical during metamorphosis, and that haemocyte functions beyond immunity are non-essential for post-embryonic development (Charroux and Royet 2009, Defaye, Evans et al. 2009, Shia, Glittenberg et al. 2009, Arefin, Kucerova et al. 2015). In contrast, ablation of embryonic haemocytes is embryonic lethal independent of control of microorganisms (Defaye, Evans et al. 2009, Shia, Glittenberg et al. 2009).

‘Haemoless’ (haemocyte-ablated) larvae and adults are more susceptible to a number of bacterial and fungal infections (Charroux and Royet 2009, Defaye, Evans et al. 2009, Shia, Glittenberg et al. 2009); however the strength of phenotype is lower than for mutations of the humoral immune system (Charroux and Royet 2009). Phagocytosis is important for the immune function of haemocytes to various pathogens, demonstrated by knock-down of phagocytic receptors, and blocking phagocytosis by injecting beads (Charroux and Royet 2009, Nehme, Quintin et al. 2011). The importance of haemocytes for the production of AMPs from the fat body is potentially stage-dependent; larvae seem dependent on haemocytes for robust AMP induction, unlike adult flies (Charroux and Royet 2009, Defaye, Evans et al. 2009, Shia, Glittenberg et al. 2009).

Studies of haemocyte functions beyond immunity show roles in phagocytosis of apoptotic cells, ECM deposition, metabolic regulation, and stem cell proliferation (Olofsson and Page 2005, Martinek, Shahab et al. 2008, Ayyaz, Li et al. 2015, Woodcock, Kierdorf et al. 2015, Shin, Cha et al. 2020). Condensation of the ventral nerve cord during embryogenesis is dependent on haemocyte migration and subsequent deposition of ECM proteins (Martinek, Shahab et al. 2008). Defects in ECM production in haemocyte-ablated embryos may contribute to embryonic lethality.

In this study, we utilised ChIP-seq data of the *Hml* gene to design an improved *Drosophila* haemocyte-specific larval and adult driver line, *Hml*^e9-P2A^-GAL4. Using *Hml*^e9-P2A^-GAL4 to drive apoptosis, we completely ablated haemocytes in the larvae. We show for the first time that haemocytes are essential for the development of adult stage flies, independent of control of the microbiota. RNA-seq data shows a striking upregulation of chitin ECM genes in the midgut of ‘haemoless’ larvae, and points to a critical role of haemocytes in regulating intestinal development.

## Results & Discussion

### *Hml*^e9-P2A^-GAL4 is a strong haemocyte-specific driver

Studies in the fly have utilised a number of different enhancer elements to achieve haemocyte-specific expression of transgenes, deriving from the *Hemese* (*He*), *eater* and *Hemolectin* (*Hml*) genes in larvae and adults and from *Peroxidasin* (*Pxn*) and *serpent (srp)* in embryos (Charroux and Royet 2009, Defaye, Evans et al. 2009, Shia, Glittenberg et al. 2009, Arefin, Kucerova et al. 2015, Csordas, Grawe et al. 2020). The most widely used haemocyte-specific driver in larvae and adults is derived from the *Hml* promoter. The first *Hml-*GAL4 driver, utilising 3kb upstream sequence of the *Hml* gene, was found to also include a second gene, *tsp68C* (Goto, Kumagai et al. 2001). Subsequently, a shorter 840bp region of the *Hml* enhancer lacking the *tsp68C* gene, *Hml*^Δ^, was found to be sufficient for haemocyte-specific expression of GAL4, and is widely used to date (Sinenko and Mathey-Prevot 2004).

To generate a stronger haemocyte-specific driver line, we optimised the *Hml* enhancer element (Fig. 1A). ChIP-seq analysis of the *Hml* gene from plasmatocytes revealed that histone H3 lysine 4 monomethylation (H3K4me1), which is typically found in enhancer regions (Calo and Wysocka 2013), extended into the *Hml* coding sequence up to exon 9 (Streeck *et al*. manuscript in preparation). We therefore included the first 9 exons of *Hml* followed by a P2A self-cleaving sequence upstream of GAL4 (*Hml*^e9-P2A^-GAL4). Transgenic flies were generated at two landing-sites, attP40 (chromosome II) and attP2 (chromosome III). To compare the expression strength of *Hml*^e9-P2A^-GAL4 with *Hml*^Δ^-GAL4 we used a *UAS-2xEGFP* reporter line (*Hml*^e9-P2A^>GFP and *Hml*^Δ^>GFP), and assayed GFP expression from isolated 3^rd^ instar larvae haemocytes by flow cytometry (Fig. 1B). *Hml*^e9-P2A^>GFP haemocytes showed ∼4-fold higher GFP expression than *Hml*^Δ^>GFP haemocytes, irrespective of the *Hml*^e9-P2A^-GAL4 landing site. This difference in expression strength was clearly observable by whole mount microscopy of 3^rd^ instar larvae and adults (Fig. 1C and Fig. S1A, B).

**Figure 1.**
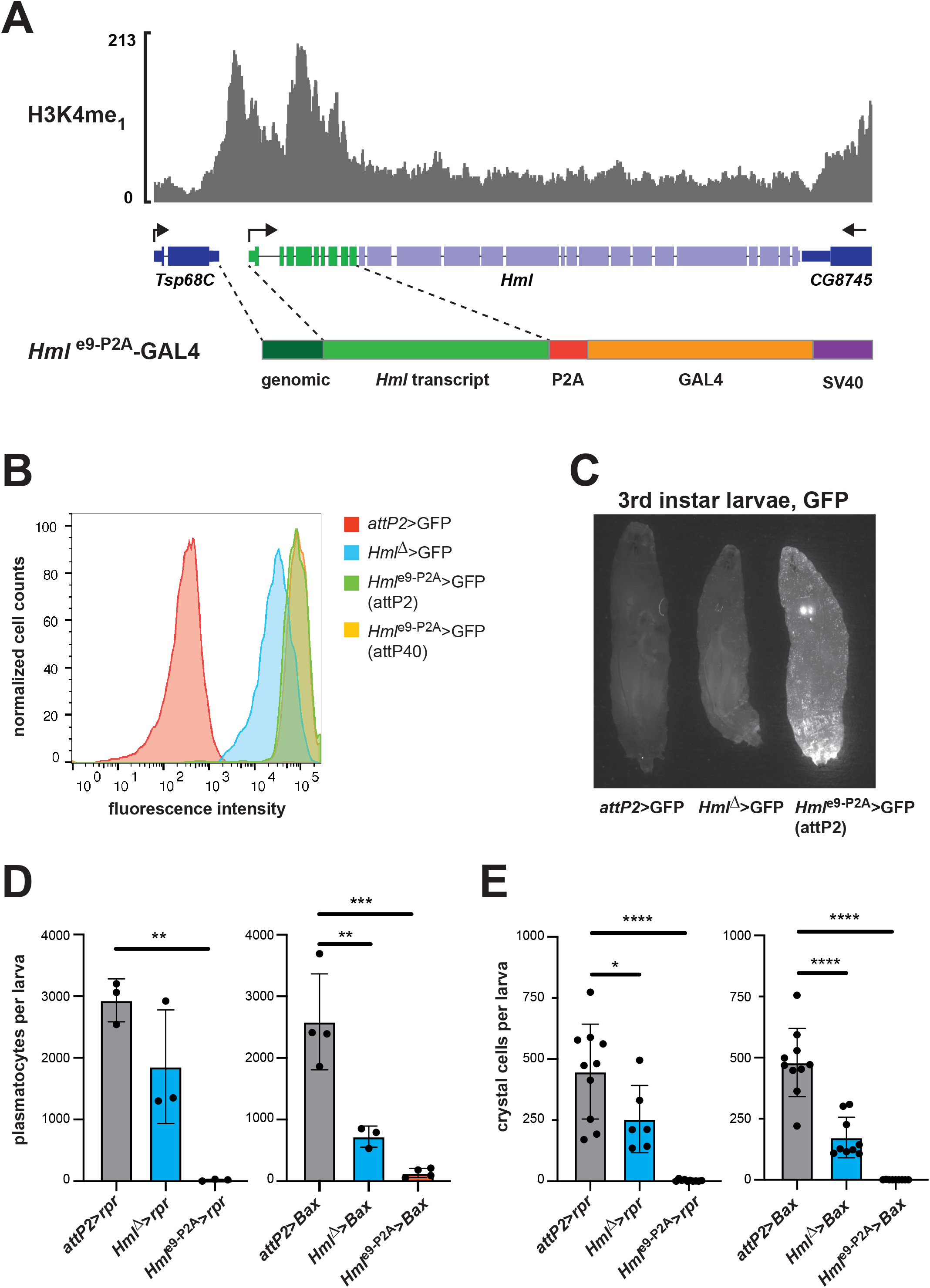
*Hml*^e9-P2A^-GAL4 is a strong haemocyte-specific driver. (A) Schematic of H3K4me1 ChIP-seq data on the *Hml* gene and the *Hml*^e9-P2A^-GAL4 construct. (B) FACS analysis of *Hml*^e9-P2A^>GFP 3^rd^ instar wandering stage larval haemocytes derived from integrations in attP40 and attP2 compared to *Hml*^*Δ*^>GFP or *attP2*>GFP (control). Representative histogram from two independent experiments. (C) Whole-mount fluorescence microscopy of 3^rd^ instar wandering stage larvae from *Hml*^e9-P2A^ >GFP, *Hml*^*Δ*^>GFP or *attP2*>GFP (control). (D, E) Plasmatocyte counts by haemocytometer (D) and crystal cell counts by whole mount microscopy (E) from 3^rd^ instar wandering stage larvae. Each dot represents average counts from 5 animals (E) or a single animal (F). One-way ANOVA analyses were performed.

Haemocyte-ablation experiments have previously been performed with *Hml*^Δ^-GAL4 driven expression of various pro-apoptotic genes, resulting in a significant reduction of plasmatocyte and crystal cell numbers (Charroux and Royet 2009, Defaye, Evans et al. 2009, Shia, Glittenberg et al. 2009, Arefin, Kucerova et al. 2015). Based on GFP reporter expression, we asked whether *Hml*^e9-P2A^-GAL4 might allow for complete ablation of haemocytes. Therefore, we expressed the *Drosophila* pro-apoptotic gene *reaper* (*rpr*) or the mouse BCL2-associated X protein gene (*Bax*) with *Hml*^Δ^-GAL4 or *Hml*^e9-P2A^-GAL4 and compared them to a control without GAL4 induction (*attP2*>*rpr*; *attP2*>*Bax*). In 3^rd^ instar *Hml*^e9-P2A^*>rpr* or *Hml*^e9-P2A^ *>Bax* larvae, we found less than 1% of plasmatocytes (Fig. 1E) and crystal cells (Fig. 1F and Fig. S1C) remaining. In comparison, driving expression with the original *Hml*^Δ^-GAL4 line resulted in reduced but detectable numbers of both blood cell type, similar to previous reports (Charroux and Royet 2009, Defaye, Evans et al. 2009, Shia, Glittenberg et al. 2009). We observed no increase in lamellocyte numbers when ablating with the *Hml*^e9-P2A^ driver, as determined by size and shape on a haemocytometer (n>30 3^rd^ instar larvae), in contrast to previous observations when ablation was performed with the *Hml*^*Δ*^ driver (Arefin, Kucerova et al. 2015). Two previous studies have observed increases in melanotic masses in ‘haemoless’ flies (Defaye, Evans et al. 2009, Arefin, Kucerova et al. 2015), although others not (Charroux and Royet 2009, Shia, Glittenberg et al. 2009). We observed no melanotic masses in *Hml*^*e9-P2A*^*>rpr* or *Hml*^*P2A*^*>Bax* larvae as determined by whole mount microscopy of 3^rd^ instar larvae (n>20).

### Haemocyte ablation with *Hml*^e9-P2A^-Gal4 is pupal lethal under germ-free conditions

Previous ablation studies using *Hml*^Δ^-GAL4 have shown a reduction in eclosion rates that were rescued when larvae were reared with antibiotics or under germ-free conditions (Charroux and Royet 2009, Defaye, Evans et al. 2009, Shia, Glittenberg et al. 2009, Arefin, Kucerova et al. 2015). This suggested a critical role for haemocytes in controlling microorganisms during metamorphosis. Given the improved ablation rate of haemocytes using *Hml*^e9-P2A^-GAL4 we revisited this observation. Both, *Hml*^e9-P2A^*>rpr* and *Hml*^Δ^>*rpr* larvae showed no decrease in pupariation rates compared to control larvae when reared at controlled density from 1^st^ larval instar (Fig. 2A). This is consistent with previous reports and suggested that larval development was not significantly affected by haemocyte ablation. We observed no gross delay in timing to pupariation between any of the genotypes. Eclosion rates of *Hml*^Δ^>*rpr* and *Hml*^Δ^>*Bax* pupae were reduced by approximately 25%, which was lower than previously reported (Charroux and Royet 2009, Defaye, Evans et al. 2009, Shia, Glittenberg et al. 2009, Arefin, Kucerova et al. 2015), but still statistically significant (Fig. 2B). Strikingly, the rates of eclosion for *Hml*^e9-P2A^*>rpr* and *Hml*^e9-P2A^*>Bax* pupae dropped to 0% (Fig. 2B). We then reared 1^st^ instar larvae either on food containing antibiotics (5mg/mL Ampicillin, 5mg/mL Kanamycin) or under germ-free conditions. Eclosion rates of *Hml*^Δ^>*rpr* and *Hml*^Δ^>*Bax* pupae were restored to expected levels (Fig. 2C, D). In contrast, even with antibiotics or under germ-free conditions *Hml*^e9-P2A^*>rpr* and *Hml*^e9-P2A^*>Bax* pupae did not eclose (Fig. 2C, D). This pointed to an essential role for haemocytes during pupal development, independent of control of the microbiota during metamorphosis.

**Figure 2.**
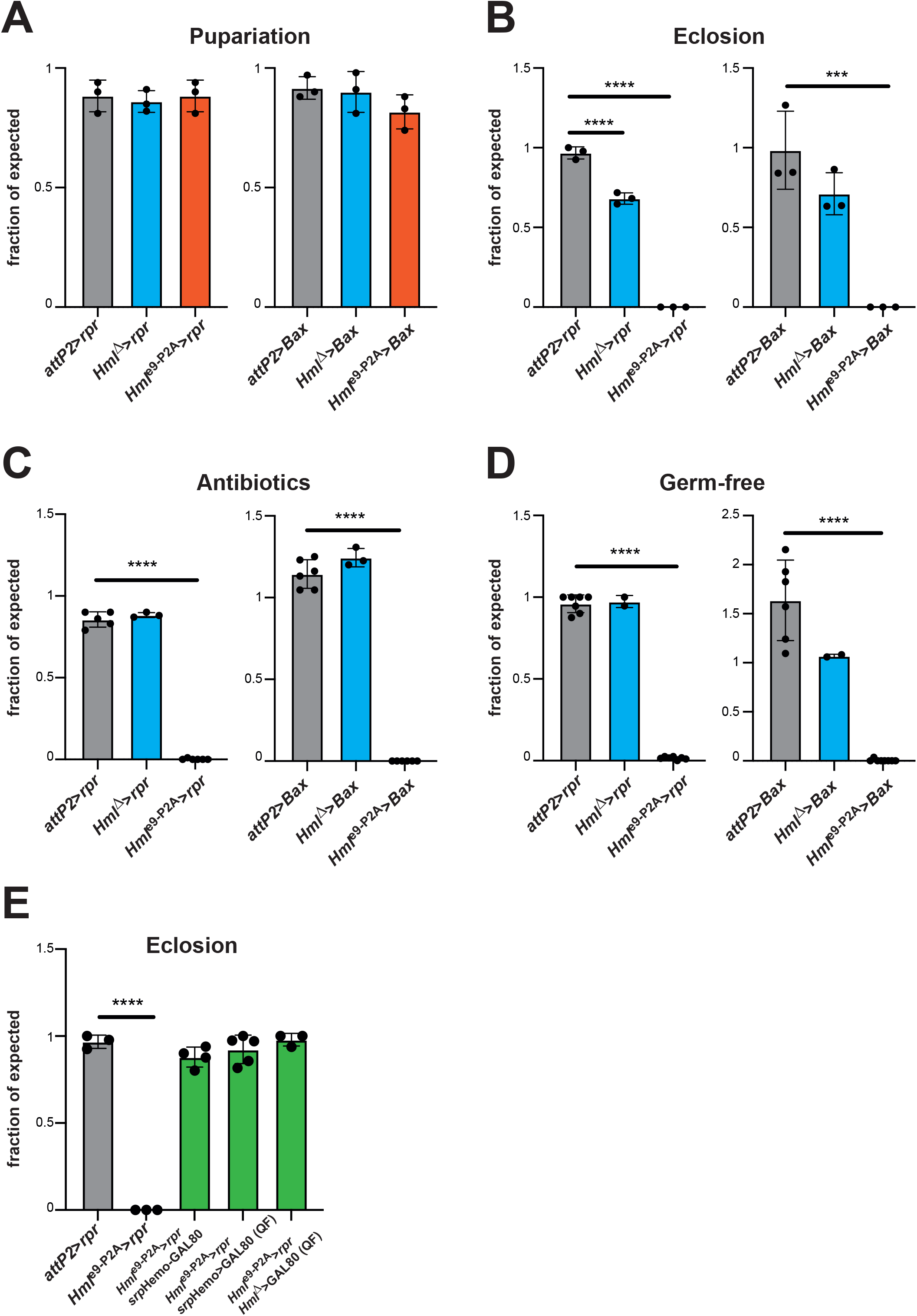
Haemocyte ablation with *Hml*^e9-P2A^-GAL4 is pupal lethal under germ-free conditions. (A) Pupariation was scored as percent of the expected number of pupae from larvae raised at controlled density (100/vial). Each dot represents an individual vial. (B) Eclosion rates were scored based on the numbers of adults as fraction of the expected number of adults. Each dot represents an individual vial. (C) Eclosion rates were scored as percent of expected adults from larvae raised at controlled density (100/vial) on standard fly food containing 5 mg/mL Ampicillin and 5 mg/mL Kanamycin. (D) Eclosion rates were scored as percent of expected adults from larvae that hatched under germ-free conditions and were raised on sterile standard fly food. (E) Genetic rescue experiments were performed supressing the activity of *Hml*^*e9-P2A*^ either by *srp*Hemo-GAL80 or through QUAS/QF mediated expression of GAL80 in *srp*Hemo>GAL80 and *Hml*^*Δ*^> GAL80 animals. Eclosion rates were scored as percent of expected adults from larvae raised at controlled density (100/vial). One-way ANOVA analyses were performed.

### Eclosion rates are rescued with haemocyte-specific expression of GAL80

In order to minimise the chance that pupal lethality in our ablation experiments was caused by off-target expression of *Hml*^e9-P2A^-GAL4, we performed genetic rescue experiments with haemocyte-specific expression of the GAL4 inhibitor, GAL80, using well established haemocyte-specific driver constructs. Since we anticipated that these experiments would be critically dependent on GAL80 expression levels we used multiple approaches. First, we used GAL80 directly regulated by a haemocyte-specific enhancer of the *serpent* (*srp*) gene (*srp*Hemo*-*GAL80), which is known to be expressed in haemocytes (Gyoergy, Roblek et al. 2018). Eclosion rates of *Hml*^e9-P2A^>*rpr, srp*Hemo*-*GAL80 pupae were restored to levels seen for the control with both *Hml*^e9-P2A^ -GAL4 lines, integrated either in attP40 or attP2 (Fig. 2E). Alternatively, we expressed GAL80 utilising the QF-QUAS system (Potter, Tasic et al. 2010). To achieve haemocyte-specific expression of a QUAS-GAL80 transgene we used *srp*Hemo-QF2 (*srp*Hemo>GAL80) or *Hml*^*Δ*^-QF2 (*Hml*^*Δ*^>GAL80). In both cases eclosion rates of *Hml*^e9-P2A^>*rpr* pupae were restored to expected levels (Fig. 2E). Pupariation was not significantly affected in any of these conditions, and the suppression of *Hml*^e9-P2A^>GFP by *srp*Hemo>GAL80 and to a lesser extent by *Hml*^*Δ*^>GAL80 was observable in whole mount microscopy of larvae and adult flies (Fig. S2A, B). Overall, this showed that pupal lethality was specifically driven by *rpr* and *Bax* expression in haemocytes, and is unlikely to result from an off-target effect.

### Haemocyte ablation leads to dysregulation of midgut expressed genes

We reasoned that effects from haemocyte ablation would potentially be detectable by transcriptional changes and therefore compared *Hml*^*e9-P2A*^*>rpr* and *attP2*>*rpr* larvae, immediately before the onset of their lethal phase by RNAseq. In this experiment downregulated transcripts would comprise both, lost haemocyte-specific transcripts and potential systemic responses; therefore, we followed two strategies to disentangle these effects. First, we compared larval RNAseq data to a plasmatocyte-specific dataset that we recently generated (Streeck *et al*. manuscript in preparation) and defined transcripts as non-plasmatocyte, shared or plasmatocyte-enriched (Fig. 3A). Second, we analysed tissue-specific enrichment of transcripts based on data available at FlyAtlas2 (Leader, Krause et al. 2018) (Fig. S3A). Known haemocyte-specific transcripts, such as *Hml, He, eater, Pxn* and *NimC1*, were detected as plasmatocyte-enriched (Fig. 3A) and were predominantly expressed in the larval carcass (Fig. S3B), likely reflecting the association of sessile haemocytes with the cuticle or lymph glands in the carcass. These transcripts were also significantly depleted in the differential expression analysis of *Hml*^*e9-P2A*^*>rpr* and *attP2*>*rpr* larvae (Fig. 3B, Fig. S3C and Table S1).

**Figure 3.**
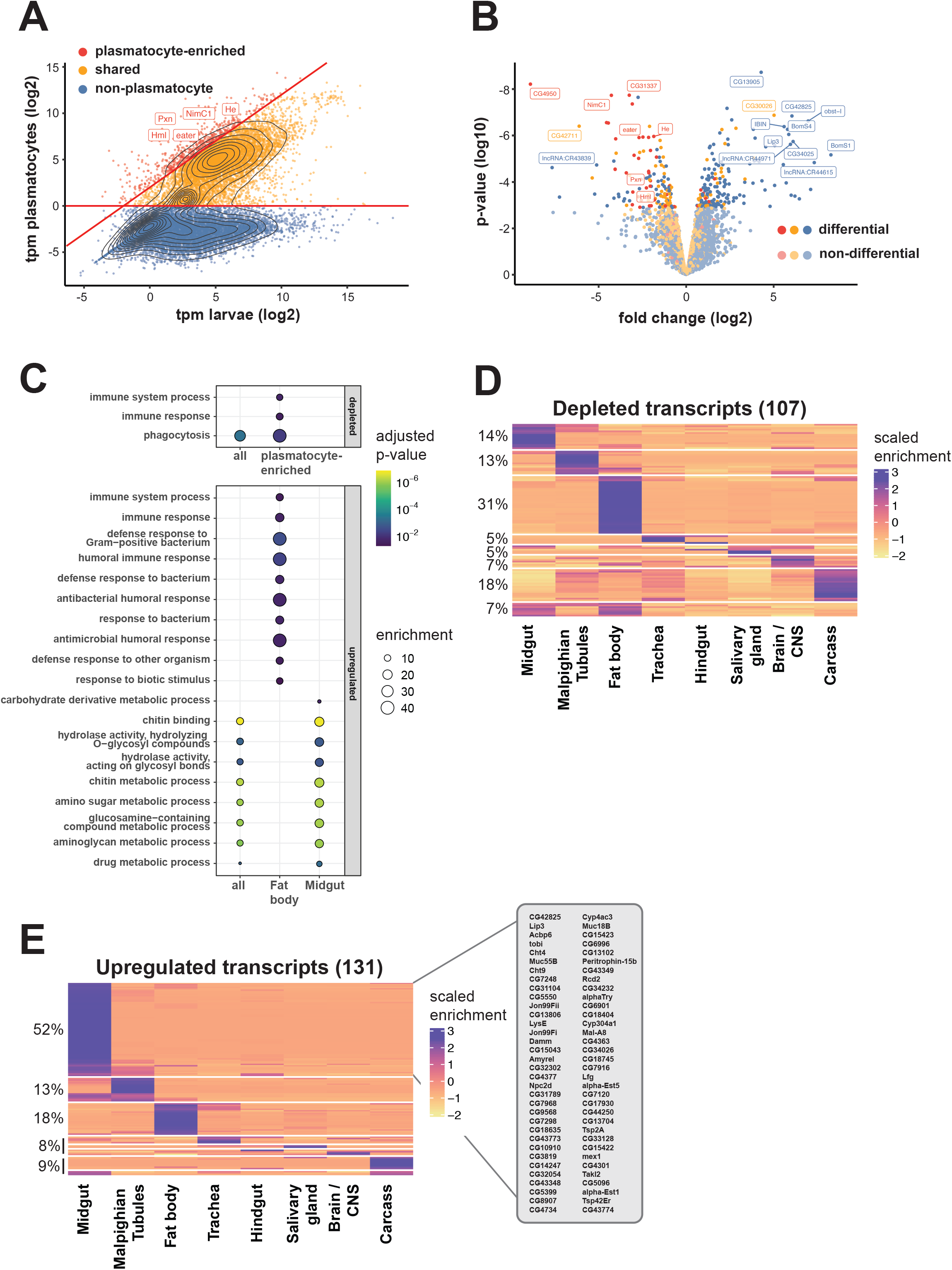
RNAseq analysis of ‘haemoless’ 3^rd^ instar wandering larvae. (A) RNAseq expression analysis comparing relative expression strength (in tags per million, tpm) of transcripts in whole larvae with expression in larval plasmatocytes (both 3rd instar wandering stage). Dots represent individual transcripts with overlaid density plot. Genes were classified as non-plasmatocyte (no or marginal expression in plasmatocytes, blue), shared (yellow) or plasmatocyte-enriched (>4 fold elevated in plasmatocytes, red). Haemocyte-specific transcripts are labeled. (B) Volcano plot illustrating differential transcriptome analysis of *Hml*^*e9-P2A*^>*rpr* versus *attP2*>*rpr* 3^rd^ instar wandering stage larvae. Dots mark log2 fold changes and log10 differential expression p-values for individual genes. Genes are colored by assignment as in with non-significantly regulated transcripts in lighter colors. The 15 most significantly regulated transcripts and haemocyte-specific transcripts are labeled. (C) Gene ontology enrichment analysis testing sets of regulated transcripts against all detected transcripts. Depleted transcripts were tested as whole set (all) or plasmatocyte-enriched subset. Upregulated transcripts were tested as whole set or as sets that show tissue specific expression (fat body, midgut). Fill color indicates p-value of enrichment and circle size shows effect size. (D, E) Heatmaps showing scaled tissue enrichment derived from FlyAtlas2 for depleted (D) or upregulated (E) protein coding transcripts. The fraction of transcripts within each k-means cluster is indicated in percent, the tissue type below the heatmap. A list of midgut specific transcripts that were upregulated in response to haemocyte ablation is shown (E). Fill color indicates scaled enrichment values.

Gene ontology (GO) analysis of depleted transcripts in ‘haemoless’ larvae revealed phagocytosis as the only significantly enriched process or function (Fig. 3C). This was likely driven by depletion of haemocyte transcripts and was maintained when plasmatocyte-enriched transcripts were analysed alone (Fig. 3C). The entire group of depleted transcripts comprised an increased fraction of fat body-(31%) and carcass-enriched (18%) transcripts relative to the control group of all genes (17% and 6% respectively) (Fig. 3D and Fig. S3A). However, only a minority of fat body-(8/34) and carcass-enriched (4/19) transcripts were non-plasmatocyte, therefore we cannot conclude that this was a tissue-specific effect of haemocyte ablation. In summary, we did not detect a clear systemic response based on downregulation of tissue-specific transcription in ‘haemoless’ larvae.

In contrast, we found upregulated transcripts primarily comprised non-plasmatocyte transcripts (131/170), indicating a systemic response to haemocyte ablation (Fig. 3B). A previous report showed that haemocyte ablation triggers a pro-inflammatory basal state (Arefin, Kucerova et al. 2015). Consistent with this, we found enrichment of immune-related GO terms within upregulated fat body expressed transcripts (Fig. 3C). The majority of upregulated genes, however, were expressed in the midgut (Fig. 3E and Fig S3E). GO analysis showed a strong enrichment of genes associated with chitin metabolism in specifically midgut-enriched genes (Fig. 3C). No other significant GO term enrichment was found for the entire set of upregulated genes. Genes regulated in the midgut included factors with chitin binding activity like *obstructor-I* (*obst-I), Mucin 55B* (*Muc55B*), and proteins with hydrolase activity such as the chitinases, *Cht4* and *Cht9* (Fig. 3C, E). These genes are critical for chitin-associated ECM production and remodelling (Pesch, Riedel et al. 2016). We speculate that the loss of haemocyte function may causes a compensatory increase in chitin-based ECM production, potentially to increase barrier function at the peritrophic membrane. Interestingly, haemocytes are found within the basal lamina of the midgut and are critical for intestinal stem cell (ISC) proliferation in response to infection (Ayyaz, Li et al. 2015). Haemocyte ablation may therefore also affect ISC maintenance, causing direct developmental defects in the gut. It would be interesting to address whether dysregulation of the midgut is a direct consequence of ablated haemocyte functions, since a reduction in midgut barrier function could explain why lethal phenotypes in partial ablation experiments were rescued under germ-free conditions, and at the same time why we found a developmental requirement of haemocytes in our complete ablation model.

Taken together, we have shown an essential role for haemocytes in post-embryonic development beyond the control of microorganisms. This new model could provide exciting insights into the requirement of haemocytes in tissue development, beyond their essential role in the immune system.

## Materials & Methods

### *Drosophila melanogaster* Strains

Fly strains obtained from Bloomington Stock Centre were *Hml*^*Δ*^*-*GAL4 (30139), *UAS-2xEGFP* (6874), *UAS-rpr* II (5824), *UAS-rpr* X (5823), *srp*Hemo*-*QF2 (78365), *srp*Hemo*-* GAL80 (78366), *Hml*^*Δ*^*-*QF2 (66468), *QUAS-*GAL80 (51590) and attP2 (8622). The *P{UAS-Bax*.*G}* integration on chromosome II (*UAS-Bax*) was a gift from Carla Saleh (Department of Virology, Institute Pasteur) and balanced with CyO, *P{ActGFP}JMR1*. The *Hml*^e9-P2A^-GAL4 lines were constructed in this study.

### Generation of *Hml*^e9-P2A^*-*GAL4 Flies

A 3477bp fragment was synthesised (Eurofins) containing 840bp upstream of the *Hml* transcription start site and the *Hml* transcript up to the end of exon 9 (bp13845367 – bp13848766, dm6), directly followed by an *Avr*II site, a P2A translation skip sequence, and a *Xba*I site. The *Xba*I site was used to insert the GAL4 coding sequence from pGawB followed by a SV40 3’UTR from pUASt. The construct was assembled in a backbone derived from pDESTR3R4-ϕ*C31att*B (Gunesdogan, Jackle et al. 2010), containing a *w*^+mc^ transformation marker and attB integration sequence. Transgenic lines were generated in *P{CaryP}attP40* and *P{CaryP}attP2* by Rainbow Transgenics. Inc (Camarillo, USA). The resulting *P{Hml-GAL4*.*e9-P2A}attP40* and *P{Hml-GAL4*.*e9-P2A}attP2* integrations were crossed out to remove integrase transgenes before further use.

### Haemocyte Ablation

*Hml*^Δ^-GAL4, *Hml*^e9-P2A^-GAL4 in attP2 or control *P{CaryP}attP2* males were crossed to either UAS-*rpr* or UAS-*Bax* virgin females. For the homozygous lethal UAS-*Bax* integration we used the absence of CyO, *P{ActGFP}JMR1* balancer chromosome to identify and score *Bax* expressing animals. Crosses were put in collection cages containing apple juice agar plates at 25°C in a 12h light/dark cycling incubator. 1^st^ instar larvae were picked from apple agar plates and placed at 100 larvae/15mL vial containing standard fly food. Larvae were reared at 25°C in a 12h light/dark cycling incubator until further experimentation.

### Pupariation/Eclosion Assays

Pupae were counted and scored from haemocyte ablation experiments as fraction of expected. For experiments involving *UAS-rpr* the score was based on pupae/vial with 100 larvae per vial. For experiments involving the homozygous lethal UAS*-Bax* integration, the score was based on the ratio between GFP positive (with balancer chromosome) and negative larvae. Similarly, eclosed adults were scored as fraction of expected adults/pupae per vial for crosses with *UAS-rpr* and by the ratio of Cy^-^ (with balancer chromosome) versus Cy^-^ flies for crosses with *UAS-Bax*. Vials were left 1 week longer to check for late eclosing adults. For animals reared with antibiotics, 5mg/mL Ampicillin + 5mg/mL Kanamycin was added to the standard fly food.

### Germ-free Flies

Crosses were set-up in collections with apple agar plates for 6hr at 25°C. Embryos were collected with PBS + 0.01% Triton-X (PBS-T) and transferred to a 100 μm cell strainer. Embryos were washed in twice in PBS-T, then placed in 70% ethanol for 5 mins. Embryos were dechorionated in 50:50 Clorax:Water (2.5% HOCl, final concentration), for 2 mins. Embryos were further washed in sterile PBS-T and pipetted into sterile standard fly food. Animals were reared at 25°C in a 12 h light/dark cycling incubator. To test for the absence of microbiota, single hatched animals were collected and crushed under sterile conditions with a pestle in 200 μL PBS. The solution was spread on YPD plates and grown at 25°C for 24 h or longer and checked for sterility.

### Genetic Rescue Experiments

Crosses were set up for the 3 different rescue strategies leading to following genotypes:

1. *srp*Hemo-GAL80 rescue: + */ UAS*-*rpr* ; *P{Hml-GAL4*.*e9-P2A}attP2 / srp*Hemo*-*GAL80
2. *srp*Hemo-QF2 rescue: *P{Hml-GAL4*.*e9-P2A}attP40 / UAS*-*rpr* ; *srp*Hemo*-*QF2 */ QUAS-*GAL80
3. *Hml*^*Δ*^*-*QF2 rescue: *Hml*^*Δ*^*-*QF2 */ UAS*-*rpr* ; *P{Hml-GAL4*.*e9-P2A}attP2 / QUAS-*GAL80

Larvae were picked and survival was scored as described above.

### Haemocyte Extraction

3^rd^ instar wandering stage larvae were collected and extensively washed under running water in 100 μm cell strainers to remove debris. Larvae were then washed in 70% ethanol for 5 mins. Sessile haemocytes were dislodged by extensively rubbing the larvae with a paint brush. Larval cuticles were ripped open from posterior to anterior using fine forceps and haemocytes bled out.

### Haemocyte Quantification

Plasmatocytes from five 3^rd^ instar larvae were bled into 20μL PBS + EDTA + protease inhibitor cocktail on parafilm and counted on a haemocytometer. For crystal cell quantification 3^rd^ instar larvae were picked and washed in PBS and then heat-shocked at 65°C for 10 mins which causes the crystal cells to melanise and turn black. For each larvae a dorsal and ventral image was taken using a Leica M205 stereo microscope. Crystal cells were counted manually.

### Plasmatocyte Fluorescence-Activated Cell Sorting Analysis

In total of 50 3^rd^ instar larvae were bled into the lid of an 1.5 mL Eppendorf tube containing 200 μL Schneider’s media w/o bicarbonate, pH 7.4 (Sigma) + 1/250 protease inhibitor cocktail (Sigma). Carcasses were removed and the cell solution was transferred to a 1.5 mL Eppendorf tube containing 800 μL fresh media and then passed through at 70 μm Flowmi tip filter (Sigma). Cells were analysed on a MACSQuant Analyser.

### Plasmatocyte RNAseq

OreR larvae were raised at controlled density as described above. Plasmatocytes were extracted as described above into complete media in tissue-culture treated dishes (Schneider’s medium with 10% FCS and 10 mM N-Acetyl-L-Cysteine). 80 larvae were bled for each sample. The larval carcasses were then removed and the plasmatocytes were allowed to attach for 10-15 minutes. Afterwards, plasmatocytes were washed 4 times with PBS and lysed in 900 μL TRIzol. Samples were moved to fresh prespun phase lock heavy tubes. 250 μL chloroform was added to each sample, mixed thoroughly and centrifuged (12000 g, room temperature, 15 minutes). The upper aqueous phase was then moved to a fresh DNA LoBind tube, and mixed with 550 μL isopropanol and 1 μL glycogen (20 mg/mL, RNAse free). Samples were mixed by inverting and incubated for 30 minutes in the freezer at -20°C. Samples were centrifuged (16000 g, 4°C, 10 minutes) and the supernatant was removed carefully without disrupting the pellet. The pellet was resuspended in 100 μL ultra-pure water with 300 mM sodium acetate and 1 μL glycogen (20 mg/mL, RNAse free). 300 μL EtOH was added and the sample was incubated for 20 minutes at -20°C, then centrifuged (16000 g, 4°C, 10 minutes) and the supernatant was discarded. The pellet was washed 2 times by adding 1 mL of 70% EtOH (prepared with ultra-pure water), each time spinning down the pellet (16000 g, 4°C, 3 minutes). Afterwards all supernatant was drained and the pellet was dried until no liquid was visible. The pellet was then resuspended in 15 μL of ultra-pure water and stored at -80°C. Libraries from samples were generated and sequenced at the Max Planck Genome Centre in Cologne.

### Larval RNAseq

*Hml*^e9-P2A^-GAL4 in attP2 or control *P{CaryP}attP2* males were crossed to *UAS-rpr* X virgins and larvae were raised as described above. 10 male larvae were collected at 3^rd^ instar wandering stage per sample across independent replicates and snap-frozen in liquid nitrogen. Larvae were transferred to Lysing Matrix E homogenization tubes with 1mL of TRIzol and ruptured on high settings in a FastPrep tissue homogenizer (MP Biomedicals). The supernatant was transferred to a fresh tube and spun down for 2 minutes at max speed. 800 μL of the TRIzol sample was transferred to a fresh prespun phase lock heavy tube (5PRIME) and 200μl Chloroform was added. Phases were separated by spinning at 12000 g, 15 min, 4°C. The upper aqueous phase was transferred to a fresh tube and mixed with 500 μL isopropanol. The resulting mix was spun at 20000 g, 15 min, 4°C. All supernatant was then drained and the pellet resuspended in 30 μL DNAse solution (Ambicon, final conc. 0.2 U/μL) and incubated for 1 h at 37°C. RNA was purified using the RNeasy Plus kit (Qiagen) by adding 270 μL RTL buffer and then isolated according the manufacturer’s instructions. RNA concentration was determined by Nanodrop and integrity was checked by Bioanalyzer. Libraries from samples were generated and sequenced at the Max Planck Genome Centre in Cologne.

### RNA data mapping and analysis

All sequencing data was transferred from the Max Planck Genome Centre (Cologne).

The reference genome fasta sequence file of the Berkeley Drosophila Genome Project assembly dm6 and the related gtf genome annotation file for dm6 of ensembl release 91 (dm6.91) were downloaded from ensembl (www.ensmbl.org) (Zerbino, Achuthan et al. 2018). A reference genome index was generated using dm6.91 using STAR-2.7.0e (Dobin, Davis et al. 2013) and used to map the fastq files. Quality control of RNA-seq mapping was performed using RSeQC (Zhang, Singh et al. 2021). All quality control files for FastQC, STAR mapping, and RSeQC were aggregated and visualized using MultiQC (Ewels, Magnusson et al. 2016) and all data was checked to make sure the library and sequencing was of good quality. Once a data set passed quality control, the gene level read counts were determined from bam files using the subread package (Liao, Smyth et al. 2013). The gene level read counts were then loaded into R. For PCA analysis the matrix of gene level read counts was transformed using the DESeq2 (Love, Huber et al. 2014) rlog function, from which the 1000 most variant genes were selected, PCA analysis was performed using the stats package prcomp function. For differential expression analysis, gene level read counts were processed using the edgeR package (Robinson, McCarthy et al. 2010) with the quasi-likelihood general linear model approach according to the manual. GO Term enrichment was performed using GOrilla (http://cbl-gorilla.cs.technion.ac.il/) (Eden, Navon et al. 2009) testing enrichment of regulated protein coding gene sets against all genes detected in the experiment

### Tissue enrichment analysis

All available data sets for protein coding genes detected in the RNAseq experiments were downloaded as text files from the web interface of FlyAtlas2 (http://flyatlas.gla.ac.uk/FlyAtlas2/index.html?page=home#). From these files tissue enrichments of individual transcripts were extracted, leaving out the enrichment in Garland cells since no documentation was available on how these cells were purified. The enrichment values were scaled in R and the resulting z-scores visualized as heatmaps after k-means clustering.

## Data Availabilty

RNA-seq data of hemocyte-ablated larvae and of primary plasmatocytes is available on ArrayExpress (E-MTAB-11095 and E-MTAB-10759 respectively).

## Figure Legends

**Supplementary Figure 1.**
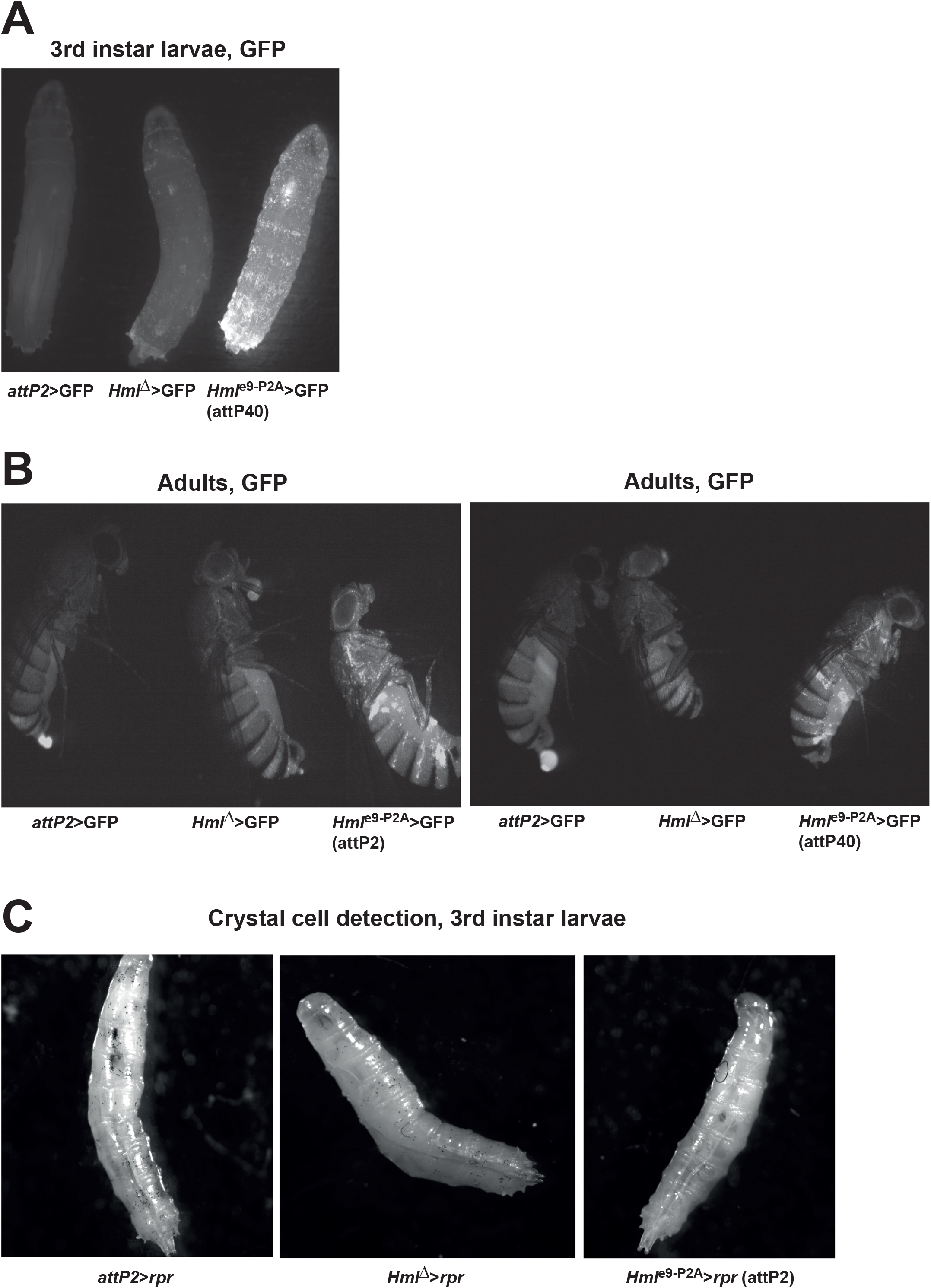
(A) Whole-mount fluorescence microscopy of *Hml*^e9-P2A^>GFP 3^rd^ instar wandering stage larvae derived from the *Hml*^e9-P2A^-GAL4 integration in attP40 compared to *Hml*^*Δ*^>GFP and *attP2*>GFP. (B, C) Whole-mount fluorescence microscopy of adult stage *Hml*^e9-P2A^>GFP animals derived from the *Hml*^e9-P2A^-GAL4 integrations in attP2 (B) or attP40 (C), compared to *Hml*^*Δ*^>GFP and *attP2*>GFP. (C) Representative images of heat-shocked larvae used for crystal cell quantification

**Supplementary Figure 2.**
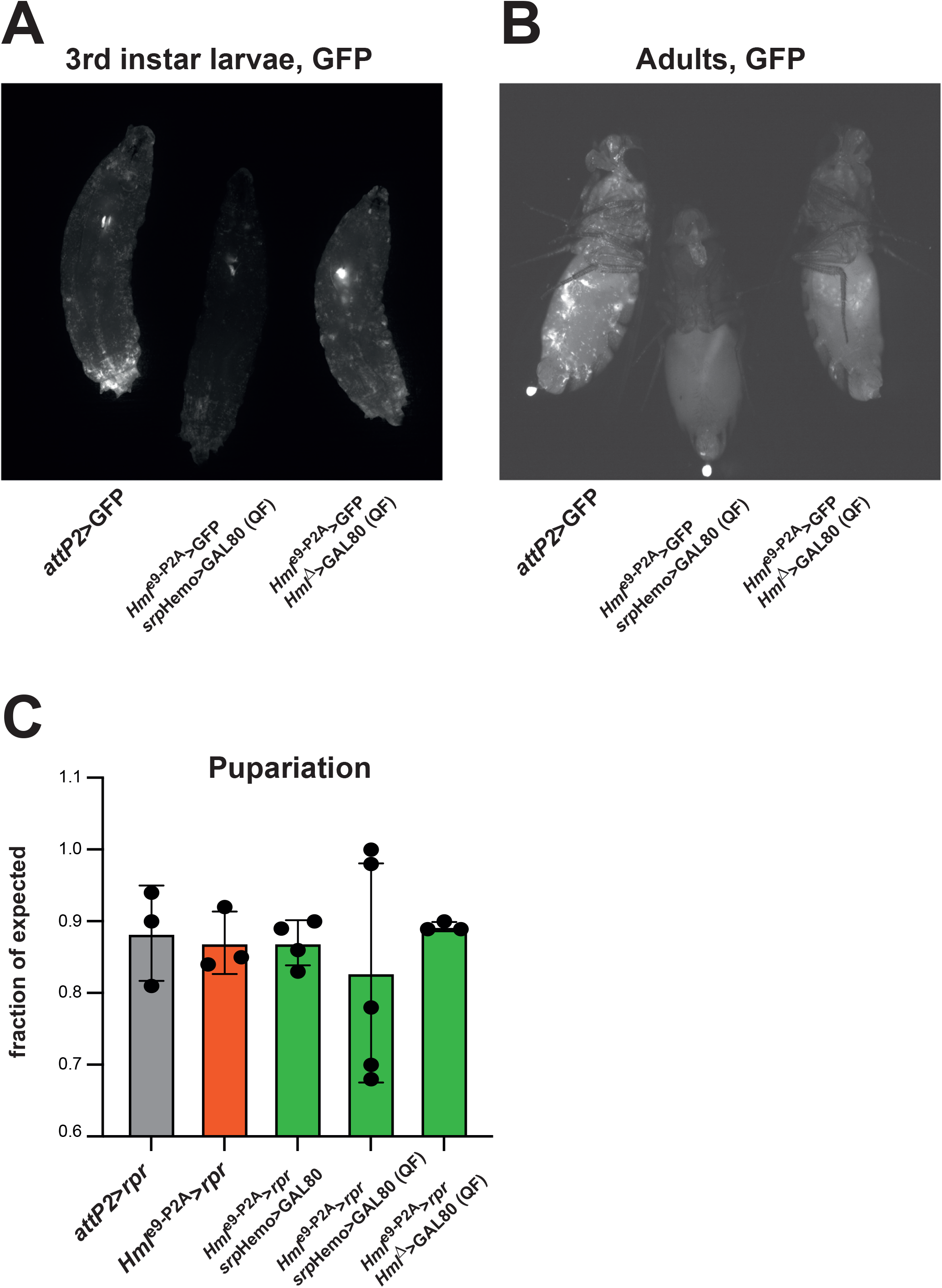
(A, B) Whole-mount fluorescence microscopy of 3^rd^ instar larvae (A) and adult flies (B) from experiments using *srp*Hemo>GAL80 and *Hml*^Δ^>GAL80 to supress GFP expression in *Hml*^e9-P2A^ >GFP animals. *srp*Hemo>GAL80 suppressed GFP expression to undetectable levels both in adults and larvae. *Hml*^Δ^>GAL80 weakened GFP expression in larvae but haemocytes were still visible. This is consistent with the idea that the *Hml*^e9-P2A^ enhancer is stronger than the *Hml*^*Δ*^ enhancer. (C) Pupariation rates were determined in genetic rescue experiments using either *srp*Hemo-GAL80 or QUAS/QF mediated expression of GAL80 in *srp*Hemo>GAL80 and *Hml*^*Δ*^> GAL80 animals to supress the activity of *Hml*^*e9-P2A*^. Rates were scored as percent of expected pupae from larvae raised at controlled density (100/vial).

**Supplementary Figure 3.**
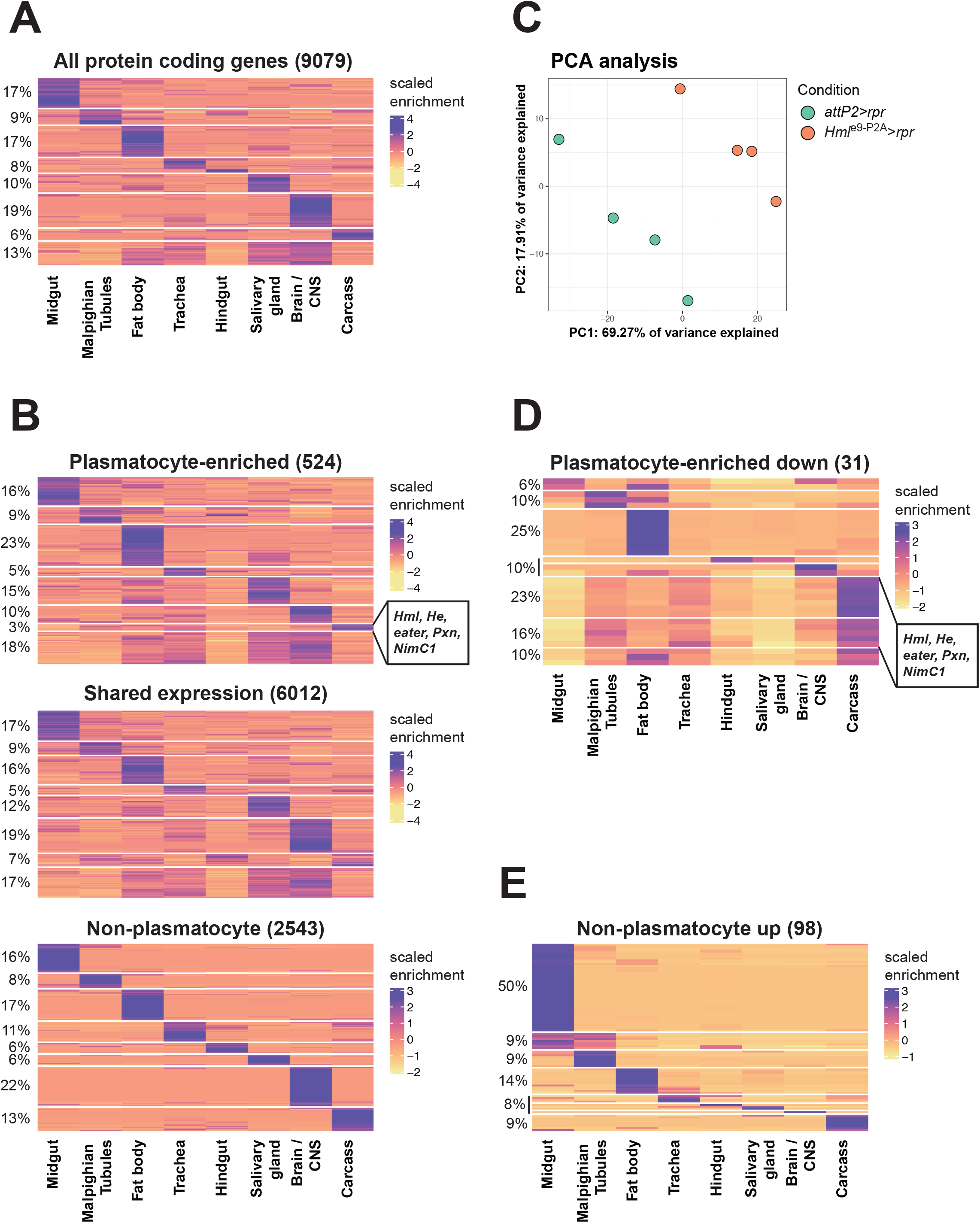
(A) Heatmap showing scaled tissue enrichment of all protein coding transcripts detected in our differential expression analysis for which data was available at FlyAtlas2. (B) Heatmaps showing tissue enrichment for subsets of transcripts shown in (A). These subsets reflect the classification into plasmatocyte-enriched transcripts, shared transcripts and non-plasmatocyte transcripts. Haemocyte specific transcripts cluster together with carcass enriched transcripts as indicated. (C) Principle component analysis (PCA) of RNAseq replicate data from *Hml*^P2A^>*rpr* or *attP2*>*rpr* 3^rd^ instar wandering stage larvae. (D, E) Heatmaps showing plasmatocyteenriched transcripts that were depleted (D), or non-plasmatocyte transcripts that were upregulated in haemocyte ablated larvae. For all heatmaps the total number of transcripts is given in brackets, the fraction of transcripts within each k-means cluster annotated as percent values and the tissue type indicated below the heatmap. Fill color indicates scaled enrichment values.

**Supplementary Table 1.**

RNAseq data from *Hml*^e9-P2A^>*rpr* compared to *attP2*>*rpr* 3^rd^ instar wandering stage larvae. Genes were classified as non-plasmatocyte (no or marginal expression in plasmatocytes), shared or plasmatocyte-enriched (>4 fold elevated in plasmatocytes).

